# Crossmodal Interaction of Flashes and Beeps Across Time and Number Follows Bayesian Causal Inference

**DOI:** 10.1101/2025.03.13.643161

**Authors:** Haocheng Zhu, Yiyang Zhang, Ulrik Beierholm, Ladan Shams

**Affiliations:** Department of Psychology, University of California, Los Angeles, USA; Department of Psychology, Durham University, UK; Department of Bioengineering, University of California, Los Angeles, USA; Interdepartmental Neuroscience Program, University of California, Los Angeles, USA

**Keywords:** Multisensory perception, Bayesian inference, Causal inference, Computational model

## Abstract

Multisensory perception requires the brain to dynamically infer causal relationships between sensory inputs across various dimensions, such as temporal and spatial attributes. Traditionally, Bayesian Causal Inference (BCI) models have generally provided a robust framework for understanding sensory processing in unidimensional settings where stimuli across sensory modalities vary along one dimension such as spatial location, or numerosity (Samad et al., 2015). However, real-world sensory processing involves multidimensional cues, where the alignment of information across multiple dimensions influences whether the brain perceives a unified or segregated source. In an effort to investigate sensory processing in more realistic conditions, this study introduces an expanded BCI model that incorporates multidimensional information, specifically numerosity and temporal discrepancies. Using a modified sound-induced flash illusion (SiFI) paradigm with manipulated audiovisual disparities, we tested the performance of the enhanced BCI model. Results showed that integration probability decreased with increasing temporal discrepancies, and our proposed multidimensional BCI model accurately predicts multisensory perception outcomes under the entire range of stimulus conditions. This multidimensional framework extends the BCI model’s applicability, providing deeper insights into the computational mechanisms underlying multisensory processing and offering a foundation for future quantitative studies on naturalistic sensory processing.

## 1. Introduction

Our brains constantly process various types of multisensory signals in daily life, continuously making decisions about whether to integrate or segregate this information (Körding et al., 2007; Shams et al., 2005; Shams & Beierholm, 2008; Stein & Stanford, 2008). For instance, when dining in a restaurant, to better understand what someone is saying, the perceptual system integrates both their voice and lip movements. In this process, it is essential to accurately infer whether the auditory and visual information originate from the same source.

Bayesian Causal Inference (BCI) which is a normative Bayesian model (Körding et al., 2007; Shams & Beierholm, 2010; 2022) has been proposed to account for human multisensory processing. In this normative framework, when processing two or more concurrent sensory inputs the nervous system relies on uncertain sensory inputs (likelihoods) and prior expectations (prior) to infer the causal structure, with the final cue combination based on the inferred structure. BCI framework has been successfully applied in several multisensory paradigms (see review by Shams & Beierholm, 2022), including ventriloquist effect (Odegaard et al., 2017; Rohe et al., 2015; Wozny & Shams, 2011), sound-induced flash illusion (Beierholm et al., 2009; Rohe et al., 2019; Odegaard et al., 2016; Odegaard et al., 2016; Zhu et al., 2024b), size-weight illusion (Peters et al., 2016), and rubber hand illusion (Marie Chancel et al., 2022; Samad et al., 2015).

Despite attempts to apply BCI models to various multisensory phenomena— including a two-dimensional model of the rubber hand illusion (Samad et al., 2015) that explicitly incorporated both spatial and temporal dimensions but did not fit human data (it was only tested in simulations)—most existing frameworks still address each dimension separately. In everyday life, however, multisensory information often unfolds across multiple dimensions (e.g., spatial, temporal, numerosity) in complex and dynamic ways. This raises a broader question: Can our perceptual system effectively integrate cross-dimensional cues to infer the causal structure of the external world? To explore this question, we propose a normative Bayesian framework that captures how the brain infers causal structures in environments with multidimensional cues. For example, when we watch a person speak, lip movements and speech are usually synchronized; if, however, there were a substantial delay (e.g., 1 second), our perceptual system would likely interpret these signals as arising from different sources rather than a single unified event. The proposed framework not only accommodates these complex facets of multisensory processing but also allows for a more generalized understanding of how laboratory findings may translate to real-world conditions, which inherently involve higher-dimensional and dynamic perceptual signals.

The sound-induced flash illusion (SiFI) (Shams et al., 2000; 2002) is one of the most prominent paradigms used to study multisensory perception (Powers et al., 2009; Rohe et al., 2019; Zhu et al., 2023; see Hirst et al., 2022 for a review). Traditionally, the SiFI has been viewed as a prime example of auditory-dominant perception: because the auditory modality tends to have lower uncertainty in the temporal domain, the visual modality is often drawn into alignment with the perceived auditory events (Shams et al., 2005). To test our multidimensional BCI model, we implemented a modified version of the SiFI paradigm that introduced discrepancies between auditory and visual signals in both the numerosity and temporal dimensions. By treating these dimensions as separate pieces of evidence, we aimed to determine whether participants could infer the causal structure of multisensory stimuli in a multidimensional context.

Qualitatively, Bayesian Causal Inference would predict that the interaction between the two modalities (and the degree of illusion) would depend on the inference of a common cause. The inference of a common cause would in turn depend on the overall consistency between the sensory inputs (here, auditory and visual) as well as the prior expectation of a common source. The overall consistency between the sensory inputs would depend on consistency across all observed features, here numerosity and timing. Therefore, moderate consistency across both dimensions may result in a similar outcome to a strong consistency in one dimension and weak consistency in another. Because multiple factors influence the inference of the causal structure as well as the inference of the sources (e.g., the number of flashes and beeps), to explore whether the human perceptual system follows Bayesian causal inference, we quantitatively compared the response distributions of each observer with that of the Bayesian Causal Inference (see section **Model Comparison**).

Our findings support the idea that the brain’s multisensory processing aligns well with multidimensional BCI, demonstrating that the perceptual system can integrate information from distinct dimensions by accounting for the varying degrees of uncertainty in each modality. These results highlight the brain’s remarkable ability to reconcile complex, multidimensional signals into a coherent representation of reality, offering new insights into the sophisticated computations underlying multisensory perception.

## 2. Methods

### 2.1 Participants

Twenty-six participants were initially recruited for this study. However, three participants were excluded for failing to meet the inclusion criteria or for performing inadequately in the unisensory conditions of the numerosity judgment task. This resulted in a final sample of 24 participants, aged 17 to 28 years (mean age = 20.5 years, SD = 2.1). All drawn from the UCLA Psychology Department’s subject pool.

The inclusion criteria required participants to have normal or corrected-to-normal vision and hearing. Additionally, they could not have a personal history of epilepsy, photosensitivity, or migraines. All participants were naive to the objectives of the study and provided informed consent through an online form prior to participation. Participants received UCLA Sona credits as part of their academic involvement in the study. The recruitment process, informed consent, and experimental procedures were reviewed and approved by the Institutional Review Board (IRB) at UCLA.

### 2.2 Apparatus and stimuli

The experiment was conducted on an Apple Mac Mini computer Model A1347 and a CRT monitor with a screen resolution of 1920 × 1080 pixels and a refresh rate of 100 Hz. This ensured consistent presentation of visual stimuli and timing accuracy across all trials. The experiment was programmed in MATLAB R2024a using Psychtoolbox, and an Eyelink 1000 eye-tracking system was integrated to ensure participants maintained fixation on a centrally presented cross throughout the task.

Visual stimuli consisted of circular white disks with a diameter of 2° of visual angle, presented for 10 ms at maximum luminance. The disks were positioned approximately 5° below the fixation cross. Auditory stimuli consisted of 10 ms tones at 3.5 kHz. The sound volume was standardized to 80 dB SPL to ensure consistent auditory input across participants. On each trial, participants were presented with one, two, or no flashes paired with one, two, or no beeps. The Inter-Stimulus Interval (ISI) between consecutive beeps or flashes was fixed at 50 ms. The first flash and first beep were presented at one of four possible SOAs: 0, 150, 300, 500 ms.

### 2.3 Procedure

The experiment was conducted in a quiet, dark lab environment. Participants sat 55–60 cm from the screen with their chins rested on a chinrest, instructed to maintain fixation on a centrally displayed cross throughout the task. Eye movements were monitored using an Eyelink 1000 device, and participants could only proceed to the next trial when their gaze was fixed on the cross.

The experiment included 20 conditions: 8 combinations of flash-beep pairings at four Stimulus Onset Asynchronies (SOAs: 0, 150, 300, and 500 ms), and 4 unisensory conditions (see Fig. 1a). The SOAs represented the temporal delay between the first beep and the first flash, with an SOA of 0 ms indicating simultaneous presentation and an SOA of 500 ms indicating a 500 ms delay. Ten repetitions of each condition, resulting in a total of 200 trials, were presented in a pseudorandom order to each participant. The inter-trial interval was 50 ms.

**Fig. 1.**
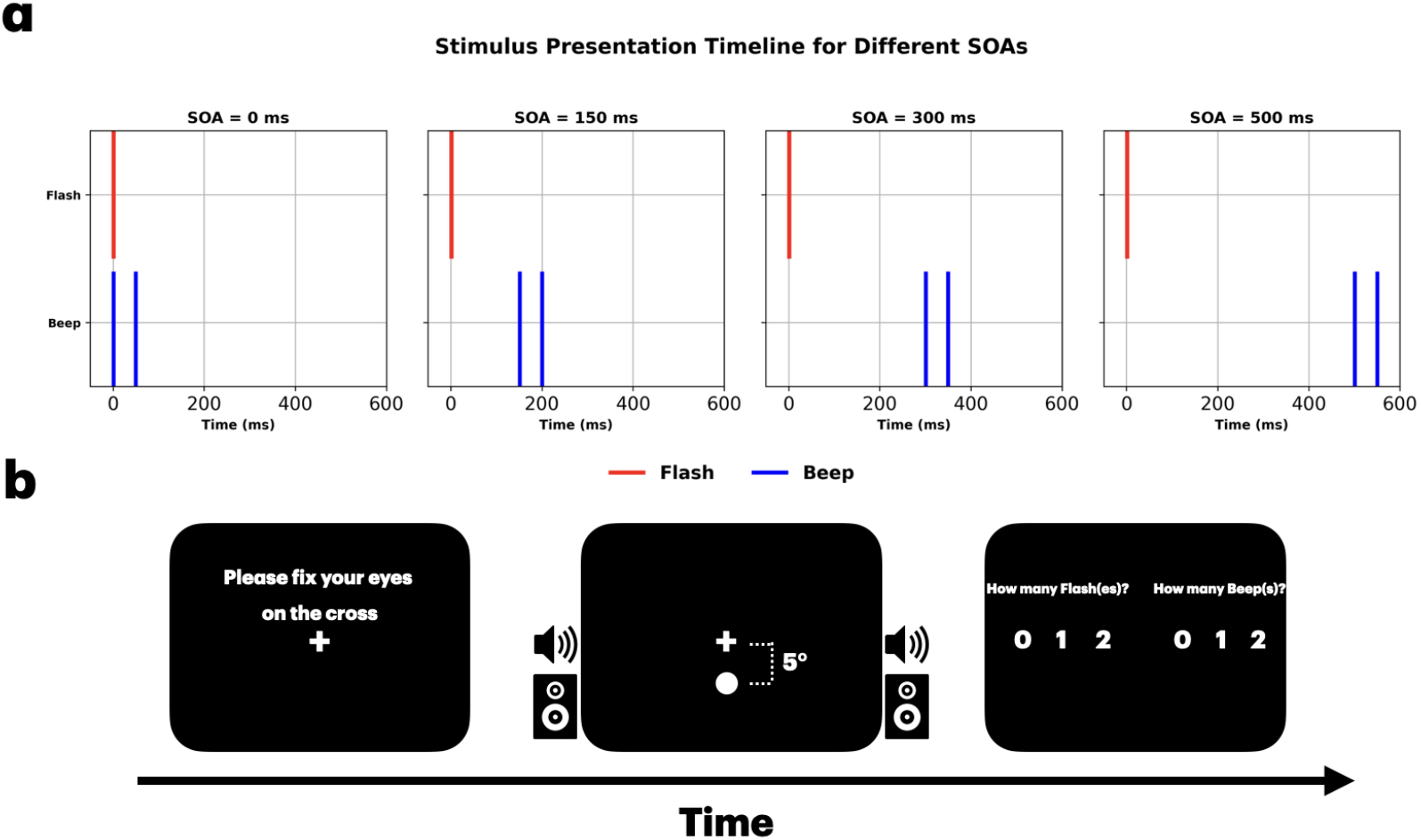
Experimental setup. (a) The stimulus timing across the four SOAs (0 ms, 150 ms, 300 ms, and 500 ms) is shown for an example flash-beep combination of 1 Flash and 2 Beeps (1F2B). Red and blue lines represent visual (flash) and auditory (beep) stimuli, respectively. (b) The trial flow: participants fixate on a central cross, then are presented with auditory-visual stimuli, followed by a prompt for their response, reporting both the number of flashes and beeps on each trial.

**Fig. 2.**
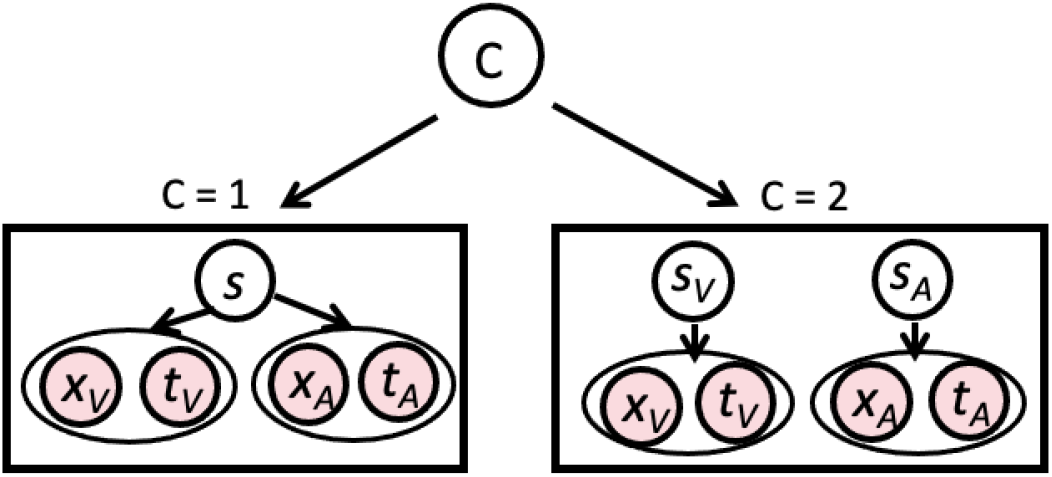
The generative model of multidimensional Bayesian Causal Inference model. Each sensory input (*A* and *V*) has two dimensions: numerosity (*x*) and time (*t*). Either both auditory and visual inputs are caused by the same source (*s*), or by two independent sources (*s*_*V*_ and *s*_*A*_).

On each trial, participants reported both the perceived number of flashes and the perceived number of beeps by pressing ‘0,’ ‘1,’ or ‘2’ (see Fig. 1b). Break screens appeared every 50 trials, allowing participants to rest and reduce fatigue. Including instructions, breaks, and trials, the experiment lasted approximately 30–40 minutes.

#### Multidimensional Bayesian causal inference model

In principle, the general implementation is based on the Bayesian causal inference model of multisensory perception (Körding et al., 2007; Wozny et al., 2010; Zhu et al., 2024b). To account for the multidimensionality, we extended the classic BCI model to encompass multiple factors. In BCI, prior expectations and current sensory information are used to infer whether sensory stimuli originate from a common cause. Here the sensory input consists of the numerosity (xV, xA; representing number of flashes and number of beeps, respectively) as well as timing of stimuli (tV, tA; representing onset of visual stream, and onset of auditory stream, respectively). The inference of the causal structure is computed according to Bayes’ rule:

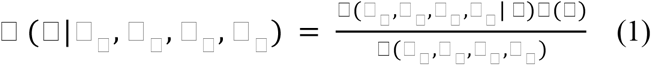

During the inference stage of perception, these two hypotheses compete to explain the sensory observations in order to estimate the perceptual variables of interest, e.g., the numerosity of the auditory event (*s*_*A*_) and the numerosity of the visual event (*s*_*V*_). The underlying causal structure of the stimuli is inferred based on the available sensory evidence and prior knowledge. Each stimulus or event *s* in the world causes a noisy sensation *x*_*i*_ of the event. We use the generative model to simulate experimental trials and subject responses by performing Monte Carlo simulations. Each sensation is modeled using the likelihood function *p*(*x*_*i*_|*s*). Trial-to-trial variability is introduced by sampling from a normal distribution around the true locations *s*_*A*_ and *s*_*V*_. This simulates the corruption of auditory and visual signals by independent Gaussian noise with standard deviation *σ*_*A*_ and *σ*_*V*_, respectively (Eq. 2-3). And the timing of auditory and visual streams are simulated by independent Gaussian noise with standard deviation *σ*_*At*_ and *σ*_*Vt*_, respectively (Eq. 4-5).

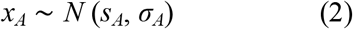

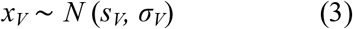

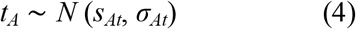

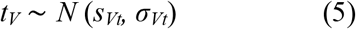

We assume that number (*x*_*V*_ and *x*_*A*_) and time (*t*_*V*_ and *t*_*A*_) are conditionally independent given □. The posterior probability of a single cause can be computed by:

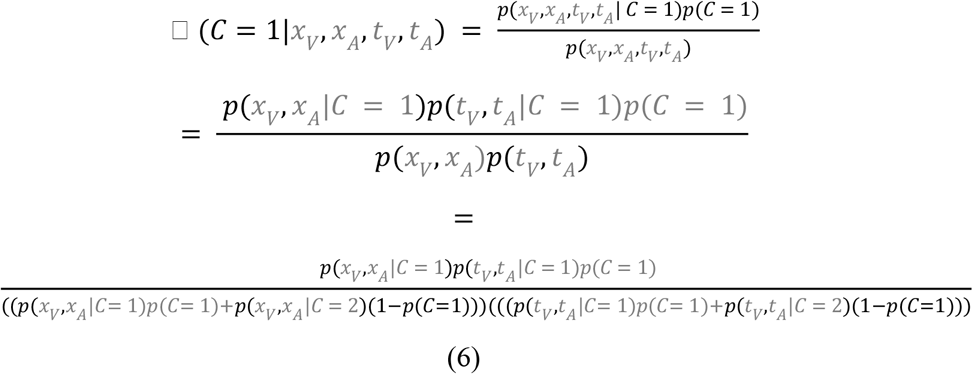

where *p(C = 1)* is the prior probability of a common cause. The likelihood of experiencing the joint sensations *x*_*A*_ and *x*_*V*_, and the timing of streams *t*_*A*_ and *t*_*V*_ given a causal structure can be found by integrating over the latent variable *s*_*i*_:

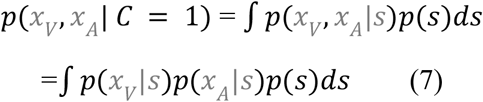

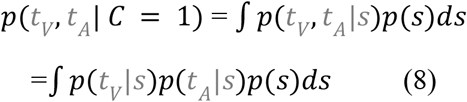

For *p(x*_*V*_, *x*_*A*_|*C= 2)*, we observe that *x*_*V*_ and *x*_*A*_ and *t*_*V*_ and *t*_*A*_ are mutually independent, allowing us to express them as the product of two separate terms:

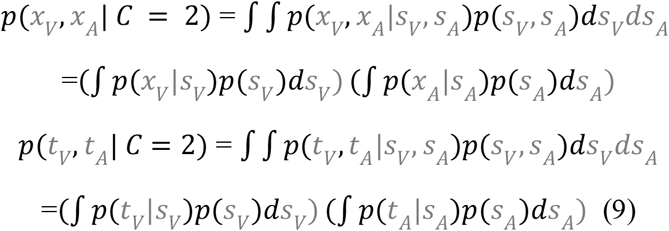

The optimal estimation of the sources within each modality is contingent on the causal structure. In the numerosity dimension, if the sensations arise from independent causes, the estimate of *each source* is computed as follows:

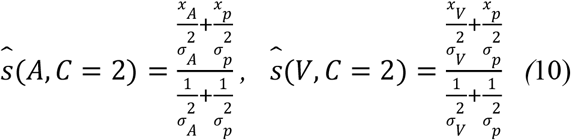

If the sensations are produced by a common cause, the estimate of *s* is computed as follows:

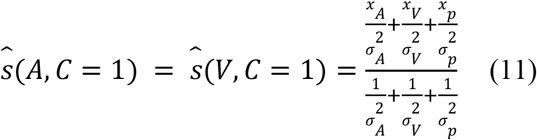

As shown in Eq. (4), the inference regarding the causal structure is probabilistic, leading to uncertainty. The optimal estimates of the sources s_V_ and s_A_ depends on the objective of the perceptual system in the specific task at hand. If the objective is to minimize the average error in the estimated source magnitudes— specifically using a sum of squared errors as the cost function—then the optimal approach is model averaging. This method incorporates the estimates from both causal structures, weighting them according to their respective probabilities (Körding et al., 2007; Wozny et al., 2010).

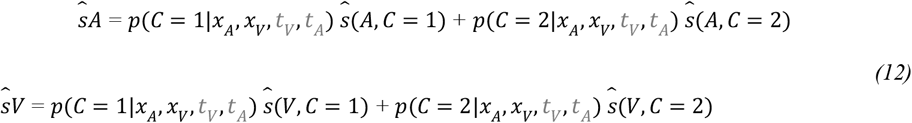

## 3. Results

We examined the perceived number of flashes and beeps as a function of discrepancy between the two modalities both in numerosity and in time. Because the accuracy of auditory perception in this task is near the ceiling in all conditions, we primarily focused on examining visual perception. We first analyzed the results from conditions where participants were exposed to varying stimulus onset asynchronies (SOAs) in the sound-induced flash illusion (SiFI) paradigm. Subsequently, we assessed how these judgments compare across conditions with different combinations of flashes and beeps. Finally, we compared participant’s responses with predictions of BCI and quantified the goodness of fit of the model.

### 3.1 Accuracy

While the experiment encompassed twenty unique conditions, our primary focus was on evaluating the effects of SOA on fusion (F2B1) and fission (F1B2) illusions. We first conducted descriptive analyses, followed by statistical evaluations to discern the significance of these effects on perceptual accuracy.

As can be seen in Fig. 3b, in 1F2B condition, visual accuracy increased substantially from 0.43 at SOA 0 ms to 0.78 at SOA 500 ms. Similarly, in the 2F1B condition, accuracy rose from 0.71 at SOA 0 ms to 0.89 at SOA 500 ms.

**Fig. 3.**
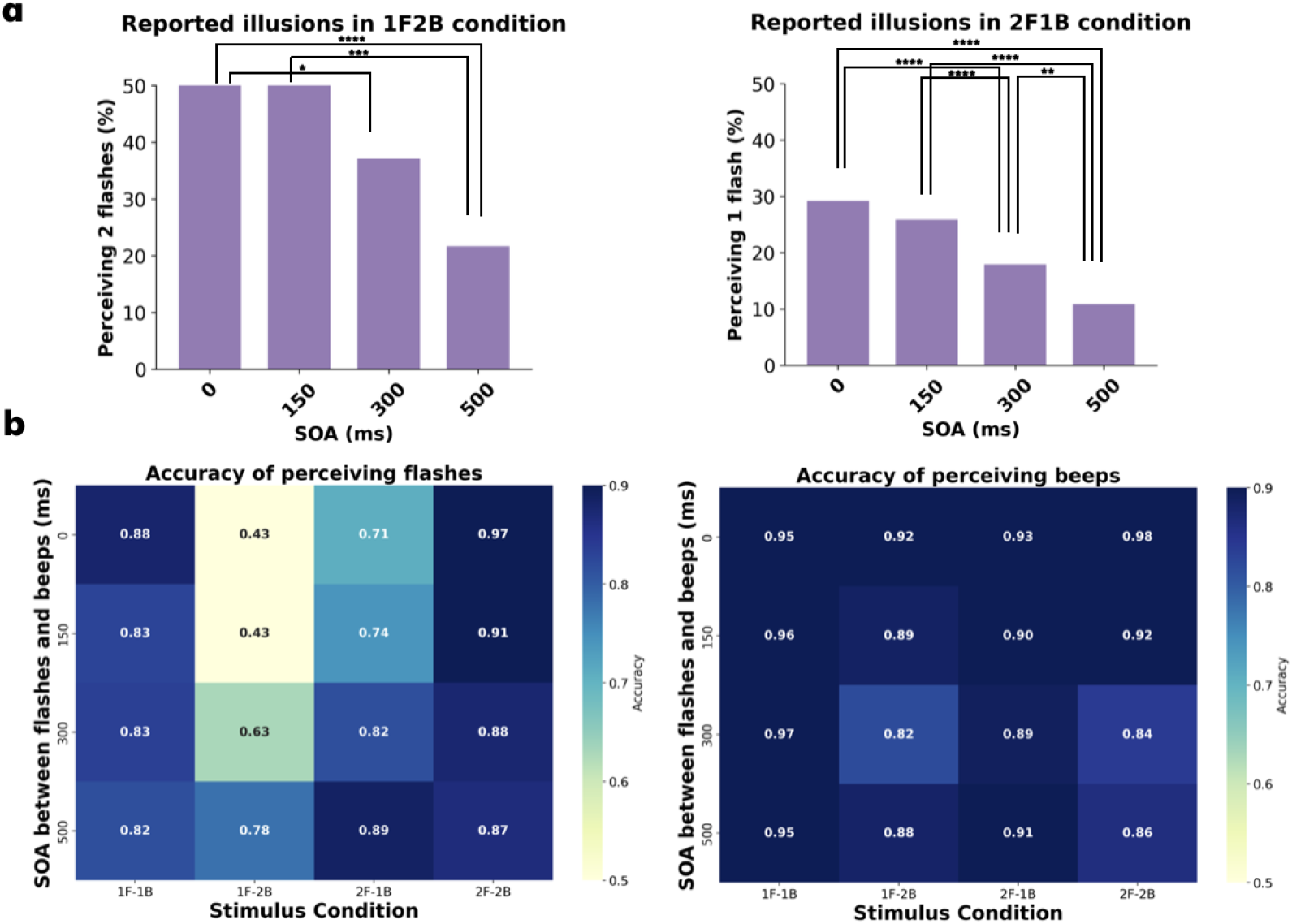
Effect of SOA on perception of Sound-induced Flash Illusion. (a) Illusion in the 1F2B (left) and 2F1B (right) stimulus conditions across varying SOAs between the stream of flashes and stream of beeps. There is a marked decrease in illusion with increasing SOA for both illusions (\*\*\*\**p* < .0001). (b) The accuracy of flash and beep response across the twenty unique experimental conditions defined by different SOA intervals and flash-beep combinations. In these figures, accuracy is plotted for each flash-beep combination (1F1B, 1F2B, 2F1B, and 2F2B) along the horizontal axis, with SOA (0, 150, 300, and 500 ms) on the vertical axis. The accuracy values are color-coded, where darker shades represent higher accuracy, and lighter shades indicate lower accuracy.

**Figure 4.**
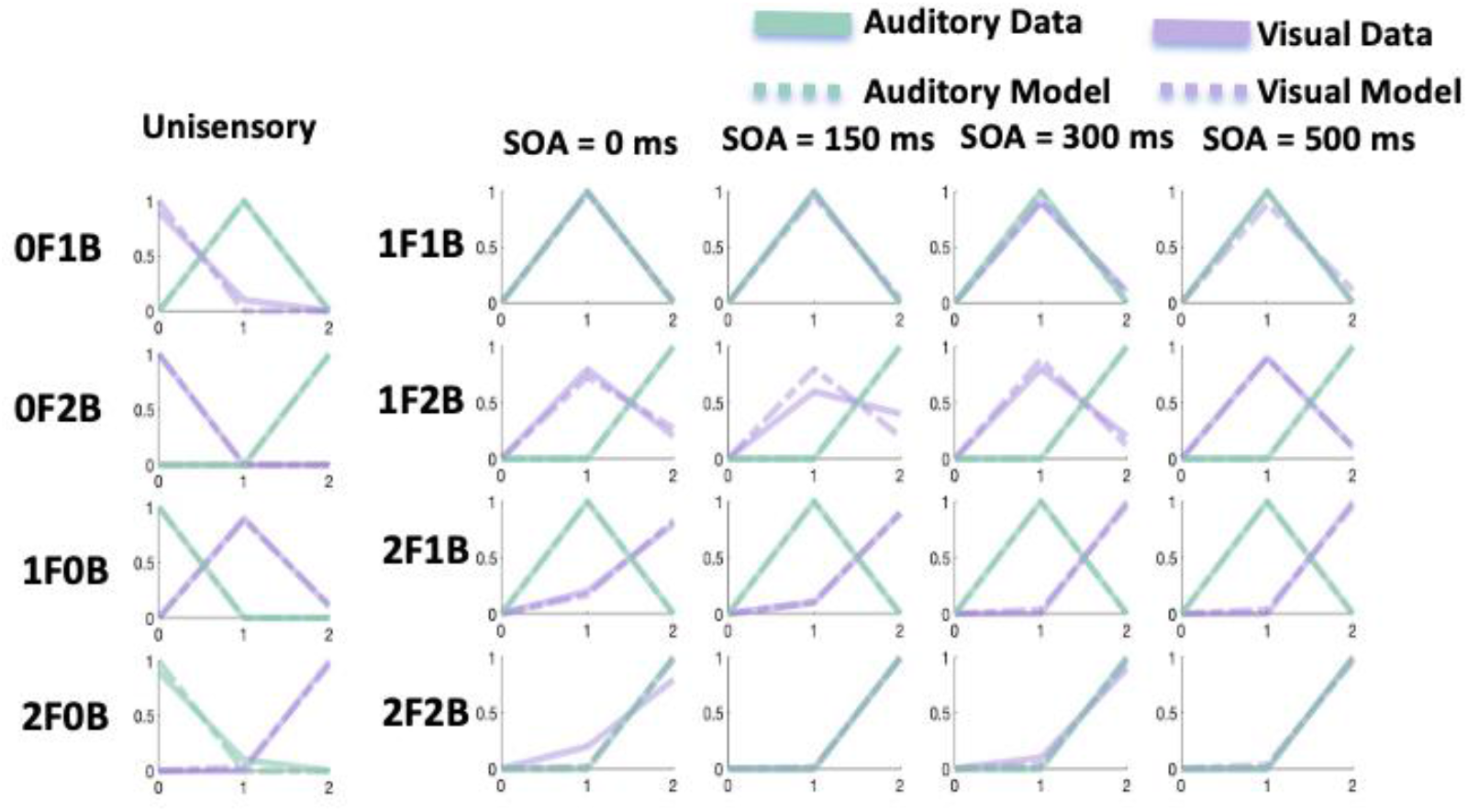
The model fitting results for a representative subject. Solid purple lines correspond to the visual response data, and green lines correspond to the auditory response data. The dashed purple lines represent the model predictions for vision, and the dashed green lines represent the model predictions for audition.

In Fig. 3c, auditory accuracy remained relatively high (> 0.8) across all conditions. In 1F2B condition, accuracy ranges from 0.92 at SOA 0 ms to 0.82 at SOA 300 ms. In the 2F1B condition, accuracy ranges from 0.89 at SOA 300 ms to 0.93 at SOA 0 ms, indicating stable perception of auditory stimuli under different SOAs.

A repeated-measures three-way ANOVA with factors SOA (4 levels), number of flashes (3 levels), and number of beeps (3 levels) revealed significant main effects of SOA. For the 1F2B condition, the analysis showed a significant main effect of SOA, F(3, 69) = 37.98, *p* < .001. Post hoc pairwise comparisons with Bonferroni correction (adjusted *α* = 0.0083) indicated significant differences between all SOA pairs except 0 ms vs. 150 ms (*t*(69) = 0.00, *p* = 1.000). Specifically, accuracy increased progressively with longer SOAs: 0 ms vs. 300 ms (*t*(69) = −5.08, *p* < .001), 0 ms vs. 500 ms (*t*(69) = −8.97, *p* < .001), 150 ms vs. 300 ms (*t*(69) = −5.08, *p* < .001), 150 ms vs. 500 ms (*t*(69) = −8.97, *p* < .001), and 300 ms vs. 500 ms (*t*(69) = −3.89, *p* = .0014).

For the 2F1B condition, the main effect of SOA was also significant, *F*(3, 69) = 10.09, *p* < .001. Post hoc tests revealed significant differences only for 0 ms vs. 500 ms (*t*(69) = −5.03, *p* < .001) and 150 ms vs. 500 ms (*t*(69) = −4.09, *p* = .0007). Non-significant comparisons included 0 ms vs. 150 ms (*t*(69) = −0.94, *p* = 1.000), 0 ms vs. 300 ms (*t*(69) = −3.04, *p* = .020), 150 ms vs. 300 ms (*t*(69) = −2.11, *p* = .234), and 300 ms vs. 500 ms (*t*(69) = −1.99, *p* = .305).

The lack of significant differences between 0 ms and 150 ms across both conditions (p = 1.000) suggests minimal perceptual integration changes at shorter SOAs (≤150 ms). The progressive accuracy improvements at longer SOAs (e.g., 500 ms) may reflect temporal stabilization of sensory processing. The stringent Bonferroni correction suppressed marginally significant effects (e.g., uncorrected *p* = .020 for 0 ms vs. 300 ms in the 2F1B condition), highlighting the need to consider effect sizes for nuanced interpretation. All reported t-tests used residual degrees of freedom (*df* = 69) derived from the repeated-measures design.

In summary, these analyses reveal that both SOA and condition significantly affect perceptual accuracy, with shorter SOAs enhancing fusion and fission illusion, and longer SOAs generally improving flash perception accuracy. This interaction underscores the context-dependent nature of temporal separation effects on perceptual accuracy, whereby SOA modulates the perception of visual stimuli based on flash-beep pairing conditions.

### 3.2 Modeling Results

#### 3.2.1 BCI Model

Because we lacked participant report data on the timing of visual and auditory stimuli, we chose to fix the corresponding parameters (*σ*_*At*_ = 40 and *σ*_*Vt*_ = 60) based on prior research (Zhu et al., 2024b). This additional restriction simplifies the model and makes the fitting procedure more reliable. Using only five free parameters to fit 200 data points, the multidimensional BCI model demonstrated a strong ability to account for the data, achieving a mean R^2^ of 0.93 across 24 subjects. To assess the model’s performance, we estimated the parameters for each subject individually using a maximum-likelihood procedure (Table 1).

**Table 1.** Sample mean ± SE parameter estimates across participants.

| N =24 | Likelihood |  |  | Prior |  |
| --- | --- | --- | --- | --- | --- |
| | $\sigma_V$ | $\sigma_A$ | $p_{common}$ | $\sigma_p$ | $\mu_p$ |
| | $0.63 \pm 0.07$ | $0.33 \pm 0.003$ | $0.62 \pm 0.06$ | $1.33 \pm 0.24$ | $1.43 \pm 0.12$ |

#### 3.2.2 Model Comparison

We quantitatively compared the Bayesian causal inference (BCI) model with the Bayesian forced-fusion and Maximum Likelihood Estimation (MLE) models using Bayesian Information Criterion (BIC). The BCI model demonstrated significantly superior model fit (BIC = 294.56 ± 28.58) compared to both forced-fusion (BIC = 378.12 ± 20.29) and MLE (BIC = 383.21 ± 20.48) models (ANOVA: *F*(2, 20) = 4.51, *p* = .0146; Tukey’s HSD, *p* < .05). No significant difference was found between the forced-fusion and MLE models (*p* = .98), indicating similar performance when assuming a single common sensory source. Detailed methodological information and additional analyses are provided in the supplementary material (see **Supplementary Material** for details).

## 4. Discussion

In this study, we presented visual and auditory stimuli with varying numerical and temporal discrepancies to investigate how the human brain computes multisensory integration. Our results indicate that the nervous system infers the causal structure underlying sensory inputs by integrating information across multiple dimensions, ultimately deciding whether to combine or segregate these signals in a probabilistic manner.

Multisensory perception has been widely studied, with research showing its benefits for memory (Quak et al., 2015; Matusz et al., 2017; Murray et al., 2022), perceptual learning (Shams & Seitz, 2008), and decision-making (Stein et al., 2020). In our study, increasing temporal discrepancies led to a reduction in the strength of multisensory illusions in incongruent conditions and a diminished multisensory benefit in congruent conditions, indicating a weakening of integration. While these findings highlight the importance of temporal synchrony, they do not imply that integration is determined solely by a fixed temporal binding window. In fact, the Bayesian Causal Inference (BCI) model predicts that integration is driven by overall consistency across multiple sensory dimensions. For example, if stimuli are highly consistent in three out of four dimensions (such as spatial location, numerosity, and intensity), even a weak temporal consistency might be sufficient to promote integration. It is the cumulative consistency that ultimately guides the perceptual system’s decision to fuse or segregate sensory inputs.

Despite these advances, quantifying sensory integration and segregation remains a challenge. The Bayesian Causal Inference (BCI) model provides a robust framework by suggesting that the nervous system combines prior knowledge with current sensory evidence to infer the underlying causal structure (Körding et al., 2007; Rohe et al., 2019; Wozny et al., 2010; Odegaard et al., 2015; 2017; Shams & Beierholm, 2010; 2022). This framework allows for direct predictions about whether inputs should be integrated or segregated. Behavioral studies have extensively validated the BCI model (Körding et al., 2007; Wozny et al., 2011; Peters et al., 2015; Odeegard et al., 2017; Dokka et al., 2019; Chancel & Ehrsson, 2023; see Shams & Beierholm., 2022 for a review), and recent neuroimaging findings further support the notion that neural circuits perform similar causal inference processes (Rohe et al., 2015; 2019; Dokka et al., 2019; Acerbi et al., 2018; Fang et al., 2019). However, as experimental paradigms evolve to better mimic the complexity of real-world sensory environments, there is an increasing need to extend the BCI framework to account for multidimensional stimuli. In response to this challenge, Zhu et al. (2024b) proposed a modified BCI model. Building on their work, our study tested whether this multidimensional approach could generalize across a broader range of conditions, and can quantitatively account for human perceptual data.

Our findings demonstrate that the multidimensional BCI model robustly predicts multisensory processing, capturing complex interactions with high accuracy. We propose that the brain employs parallel probabilistic processes to integrate information across various dimensions—such as temporal and numerical cues—to infer the causal structure of sensory events. Notably, a discrepancy in any one dimension (e.g., a substantial temporal mismatch) can override numerical consistency, prompting the system to treat the stimuli as originating from separate sources.

While traditional multisensory studies often involve manipulating only one dimension of the perceptual variables, real-world sensory processing is inherently multidimensional. Our findings suggest that perceptual inference is more flexible and probabilistic than previously assumed, extending beyond rigid cue combination rules. This has practical applications in fields such as human-computer interaction, virtual reality, and clinical neuroscience. For example, understanding how individuals infer causal relationships in noisy environments can improve the design of assistive technologies for sensory impairments. Finally, our work opens new avenues for future quantitative investigations of multisensory integration (Zhu et al., 2024a). By systematically manipulating temporal discrepancies and other sensory features, future studies can further elucidate the mechanisms that define the temporal binding window and govern the integration and segregation processes in naturalistic settings.

## 5. Conclusion

In the present study, we proposed a novel framework for the BCI model that incorporates information from different dimensions of perceptual stimuli. Using a behavioral approach, we manipulated the numerical and temporal discrepancies between audiovisual stimuli to evaluate the model. Our findings indicate that the multidimensional BCI model provides highly accurate predictions for multisensory perception under complex conditions. This study further supports the notion that the brain performs Bayesian causal inference during multisensory processing. Moreover, our newly proposed BCI framework offers substantial potential for enabling more quantitative investigations into complex multisensory research.

## Supporting information

model comparison

## Declaration of Competing Interest

The authors declare that they have no known competing financial interests or personal relationships that could have appeared to influence the work reported in this paper.

## Data Availability

Data will be made available on request.

## Notes

### Competing Interest Statement

The authors have declared no competing interest.

