## Supplementary material for "Crossmodal Interaction of Flashes and Beeps Across Time and Number Follows Bayesian Causal Inference": model comparison

The **Bayesian forced-fusion** model operates under the assumption that all sensory inputs originate from a single, common source. It infers this source by combining sensory evidence, represented through likelihood functions with prior expectations. Essentially, this model represents a constrained version of Bayesian causal inference, where the probability of a shared origin for the inputs is fixed at 1 (Eq. S1). When both the likelihood and prior distributions are Gaussian, and the estimation follows a maximum a posteriori (MAP) approach, the inferred source is determined as a weighted sum of the two sensory signals (*x_A_* and *x_V_*) along with the prior. The contribution of each component to the final estimate is governed by its precision (or reliability), which is mathematically characterized as the inverse of its variance.

$$\hat{s} =\frac{\frac{x_{A}}{\sigma_{A}^{2}}+\frac{x_{V}}{\sigma_{V}^{2}}+\frac{x_{p}}{\sigma_{p}^{2}}}{\frac{1}{\sigma_{A}^{2}}+\frac{1}{\sigma_{V}^{2}}+\frac{1}{\sigma_{p}^{2}}} (Eq. S1)$$

The **Maximum Likelihood Estimation** (MLE) model also assumes that all signals originate from a common source, but estimates this source by maximizing the overall likelihood of the combined sensory inputs. In this framework, each signal contributes to the final estimate based on its relative precision, resulting in a weighted average of the observations (Eq. S2). Unlike the Bayesian forced-fusion model, the MLE approach does not incorporate prior information. Instead, it formulates the fusion process purely as an optimization task, relying solely on the likelihoods (or reliabilities) of the signals. This method ensures that the final estimate corresponds to the most probable percept under the assumption of a single-source origin.

$\hat{s} =\frac{\frac{x_{A}}{\sigma_{A}^{2}}+\frac{x_{V}}{\sigma_{V}^{2}}}{\frac{1}{\sigma_{A}^{2}}+\frac{1}{\sigma_{V}^{2}}} (Eq.S2)$

To quantitatively compare the Bayesian causal inference (BCI) model with the forced-fusion and Maximum Likelihood Estimation (MLE) models, we computed the Bayesian Information Criterion (BIC) for each. Our analyses indicate that the BCI model achieves a BIC of 294.56 ± 28.58, while the forced-fusion and MLE models yield BIC values of 378.12 ± 20.29 and 383.21 ± 20.48, respectively. To statistically assess whether there are significant differences in model fits, we conducted a one-way analysis of variance (ANOVA) on the BIC values across the three models. The ANOVA revealed a significant main effect of model type on BIC values (*F*(2, 20) = 4.51, *p* = .0146), indicating that at least one model significantly differs from the others in terms of fit.

To further explore these differences, we conducted post-hoc pairwise comparisons using Tukey’s Honest Significant Difference (HSD) test. The results showed that the BCI model yielded significantly lower BIC values compared to both the forced-fusion (*p* < .05) and MLE models (*p* < .05), confirming that incorporating causal inference mechanisms leads to a superior fit. In contrast, the BIC values of the forced-fusion and MLE models did not significantly differ from each other (*p* = .98), suggesting that both models perform similarly under the assumption of a fixed single-source perceptual integration.
